# On the principles of cell decision-making: intracellular coupling improves cell responses fidelity of noisy signals

**DOI:** 10.1101/125864

**Authors:** Andreas I. Reppas, Eduard A. Jorswieck, Haralampos Hatzikirou

## Abstract

Reliability of cell fate decision-making is crucial to biological development. Until today it has not been clear what are biology’s design principles that allow for reliable cell decision making under the influence of noise. Here, we attempt to answer this question by drawing an analogy between cell decision-making and information theory. We show that coupling of intracellular signalling pathway networks makes cell phenotypic responses reliable for noisy signals. As a proof of concept, we show how cis-interaction of the Notch-Delta pathway allows for increased performance under the influence of noise. Interestingly, in this case, the coupling principle leads to an efficient energy management. Finally, our postulated principle offers a compelling argument why cellular encoding is organized in a non-linear and non-hierarchical manner.

## Intoduction

Cell decisions are responses of cell’s internal mechanisms to external signals [1]. Such mechanisms are typically termed as signal transduction pathways. Signal transduction occurs when an extracellular signalling molecule bounds to a certain receptor creating a complex. In turn, this complex of molecules triggers a biochemical chain of events inside the cell, leading to a phenotypic response [2]. These responses are required to be of high fidelity and reliability since they are related to vital organism processes, such as cellular metabolism, shape, differentiation, or mitotic activity [3].

Signal transduction pathways can be considered as input-output communication channels. Each pathway involves a number of key interacting proteins. The chain of biochemical events that involves the reception, endocytosis, activation/inactivation, nucleus internalisation, degradation or exocytosis of such a protein defines a single processing channel, in terms of encoding and decoding. The network of interactions of these coupled protein-channels defines the signal transduction channel.

Signal transduction pathways are subject to noisy perturbations. Typically, intracellular or intrinsic noise corresponds to thermal/stochastic fluctuations of the involved protein interactions. Alternatively, it may also account for any unknown and unobserved protein. Extrinsic noise lumps phenomena that contribute to stochasticity in cell-cell or cell-microenvironment communication.

In this paper, we seek answers on biology’s design principles that allow for reliable cell decision-making under the influence of noise. By means of a minimalistic communication model, we show that channel coupling can enhance the information capacity. In turn, we focus on dynamic cell fate determination processes, such as the Notch-Delta pathway, by establishing an analogy with communication theoretical concepts. We show that pathway coupling improves the reliability of responses in noisy environments which, thus, can be considered as a mechanism for robust cell-decision making.

## From information theory to cell decision-making

Cells can be viewed as decision-makers that constantly process available information to adapt to their microenvironment. In particular, cells encode this information into genetic, epigenetic, transcriptional and translational “codes” required for phenotypic responses [2]. Our main *hypothesis* is that information encoding costs energy to a cell, since it involves the synthesis, transport or modification of molecules required for cellular responses [4, 5]. Therefore, cell decision-making requires efficient information encoding and transmission according to the energy availability and the cost of information.

The question that arises is how to quantify the reliability/fidelity of cell decisions. Information theory offers tools for understanding this cellular information processing. More specifically, **channel information capacity** *C* defined as:

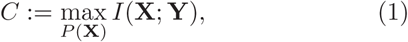
describes the maximum rate at which information can be reliably delivered over a channel. The quantity

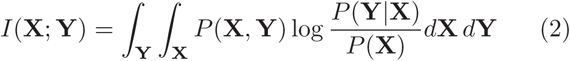
is the mutual information between the two random variables **X** and **Y** and measures the reduction in uncertainty of one variable by knowing the other [6]. The channel is described by the conditional probability function *P*(**Y|X**) with input **X**, distributed by *P*(**X**), and output **Y**. The mutual information offers a mechanism-blind treatment of cell decision-making as an input-output system, without requiring the exact knowledge of the underlying intracellular components involved in phenotypic regulation.

## A toy-model of channel coupling

The simplest model of channel coupling is the one composed of two identical binary symmetric channels (BSCs) with the corresponding inputs-outputs duplets (*x*_*i*_, *y*_*i*_), *i* = 1, 2 (Fig. 1(a)). In a BSC, the input and output are binary, i.e. (*x*_*i*_, *y*_*i*_) ∈ {0, 1}^2^ and the noise enters as an error probability, i.e.

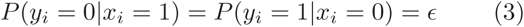
with ϵ ≤ 1/2. As a possible realisation of intracellular coupling, we assume that the output *y_j_* of one channel influences the noise ϵ of the other channel *x_i_*, where *j ≠ i* [7]. More specifically, we introduce coupling as a symmetric noise pay-off *δ* > 0. In this case, the noise of a channel, due to the presence of the other channel’s signal, becomes ϵ_*i*_ = ϵ + (− 1)^*yj*^ *δ*, with *i ≠ j* and under the sum constrain ϵ_1_ + ϵ_2_ = 2ϵ. The above model realisation ensures that when the capacity of one channel is increased the capacity of the other is decreased. The information capacity for the two coupled channel system is given by [6]:

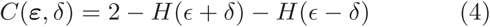
where ϵ = [ϵ + *δ, ϵ* − *δ*]. Please note that for a BSC the binary entropy function reads as *H*(*x*) = −*x* ln (*x*) − (1 − *x*) ln (1 − *x*). For small coupling strengths *δ* ≪ 1, we can obtain an explicit formula for the aforementioned capacity:

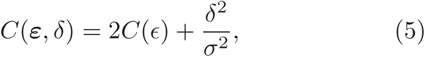
where *C*(ϵ) = 1 − *H*(ϵ) the single channel capacity and the corresponding variance *σ*^2^ = ϵ(1−ϵ). The above relationship shows readily that the coupled-channel information capacity is larger than the sum of the individual capacities which corresponds to the capacity of the uncoupled system.

**FIG. 1.**
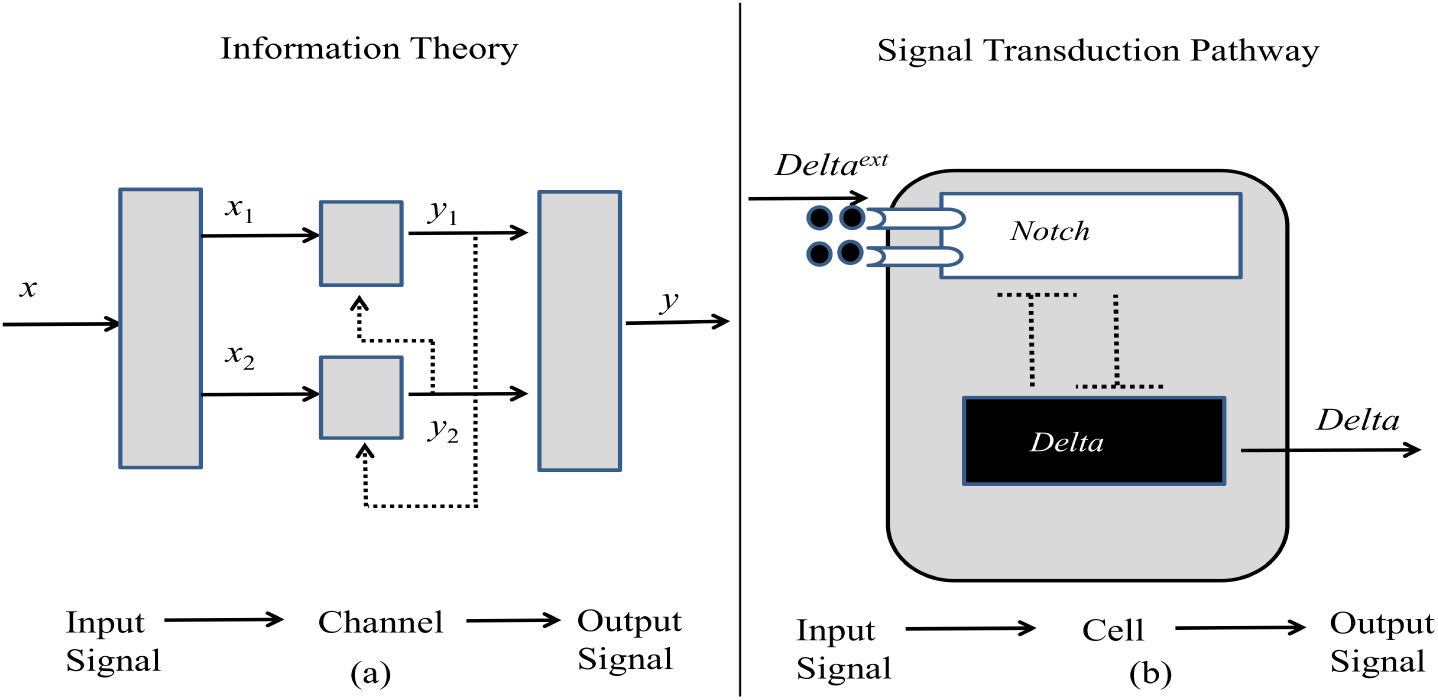
Information theory vs. Signal transduction Pathways. (a) Joint coupling (dotted arrows) of communication channels. (b) Mutual coupling (dotted inhibition arrows) of the Notch-Delta transduction channel.

Let us assume that our two-channel system operates under two different noises. This translates into a system of BSCs with high noise ϵ and one consists of channels with low noise ϵ′, with ϵ > ϵ′. Our goal is to keep the same performance by modulating the coupling strength appropriately. In the absence of coupling, we know that Δ*C*_*unc*_ = 2[*C*(ϵ′) − *C*(ϵ)] > 0. Assuming that the pair of BSCs with high noise is coupled, then its information capacity takes the form of Eq. (5). Consequently, we can find appropriate coupling strength *δ^*^ >* 0 that equalises the capacities of the coupled and uncoupled system. In this case, the coupling strength is equal to:

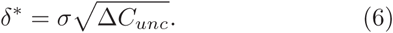
Therefore, even for high intrinsic noise channels, coupling can improve the operational fidelity and reliability of system responses. Similar results for systems of K coupled BSCs have been proven in [7]. Finally, the aforementioned result holds for Gaussian channels.

## The coupled Notch-Delta pathway

In the following, we interrogate the biological relevance of our theoretical results with respect to signal transduction pathways. In particular, we are interested in the impact of intracellular coupling in terms of response reliability. Please note that for any biological system, the idealized assumptions of BSC and symmetric noise pay-off are not valid any more. Thus, we expect a qualitative confirmation of the above results.

For a proof of principle, we focus on the Notch/Delta pathway that represents a signal transduction mechanism for cell fate determination [8]. The whole process is based on the interaction of the extracellular domain of the two transmembrane ligands Delta and Serrate on the surface of one cell with the extracellular domain of Notch receptor in a neighbouring cell. The ligand/receptor binding results in the generation of a receptor’s intracellular domain which, in turn, down-regulates Delta activity. In addition to the external activation (trans-activation) of the ligand/receptor, Notch and Delta mutually inactivate (cis-inactivation) each other inside the cell [9]. Thus, according to the above mechanisms, high levels of Delta in one cell results in high levels of the receptor’s activity in its interacting cells and vice versa. In [10], the state of high Delta/low Notch is characterised as *sender* and the complementary as *receiver*.

The Notch-Delta communication can be viewed as a two-channel network with an input received by the Notch-channel and the output produced by the Delta-channel, as in Fig. 1. Interestingly, we observe that the output of each channel is fed back to the other as an input, in the form of an inhibitory feedback. Several mathematical models of Notch-Delta signalling have been introduced to investigate the underlying mechanisms of the interaction [9, 11]. Denoting the levels of *Notch, Notch-Delta Complex* and *Delta* in a cell by *N,C* and *D*, respectively, we can write the following system of equations:

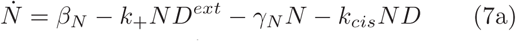

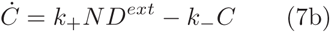

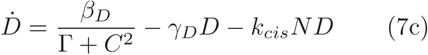
Parameter *k_+_* represent the activation rate of the complex due to the binding of Notch and Delta external (*D^ext^*). Parameters *β*_*N*_, *β*_*D*_ denote the activation rates of Notch and Delta, respectively. Parameter *k_cis_* is the rate of mutual deactivation of Notch and Delta. Parameters *γ_N_, k*_ and *γ_D_* denote the natural degradation rates of Notch, Complex and Delta, respectively. This model is similar to the one used in [9].

According to our information theoretic analogy, *D^ext^* can be viewed as a normally distributed signal, i.e. *D^ext^* ∼ *N*(*μ, σ*^2^). The internal dynamics represent the signal’s encoding and decoding into the released output Delta. The parameter *k_+_* represents the cell’s encoding strength, which can be seen as the cell’s responsiveness to the external signal, and *k_cis_* can be viewed as the coupling strength of the system. Therefore, we are particularly interested in the effect of *k_cis_* on the output fidelity. In this regard, we measure the mutual information *I*(*D^ext^; D_eq_*) of the Notch-Delta channel for variations of the cis-inhibition rate, where *D_eq_* is the steady state response of the system for a given input *D^ext^*. The mutual information was computed using the MuTE toolbox [12].

Increasing the cis-inhibition rate *k_cis_*, we observe a gain in the mutual information *I*(*D^ext^; D_eq_*), as shown in Figure 2. Assuming a high and a low noise input *D_ext_*, we observe for both a similar increase in performance by varying the coupling strength. As shown in Eq. (6), a coupling rate *k_cis_* exists that compensate for the high noise pathway performance when compared to the low noise one. We can postulate that differentiating cells can sustain the fidelity of their fate decisions by adapting their cis-inhibition rates according to the current input noise. The latter is in line with previous results indicating that cis-inhibition coupling enhances fate selection accuracy in a multicellular system [13, 14]. The aforementioned *k_cis_* effect could be easily validated in single cell experiments by measuring the mutual information for different noisy inputs.

**FIG. 2.**
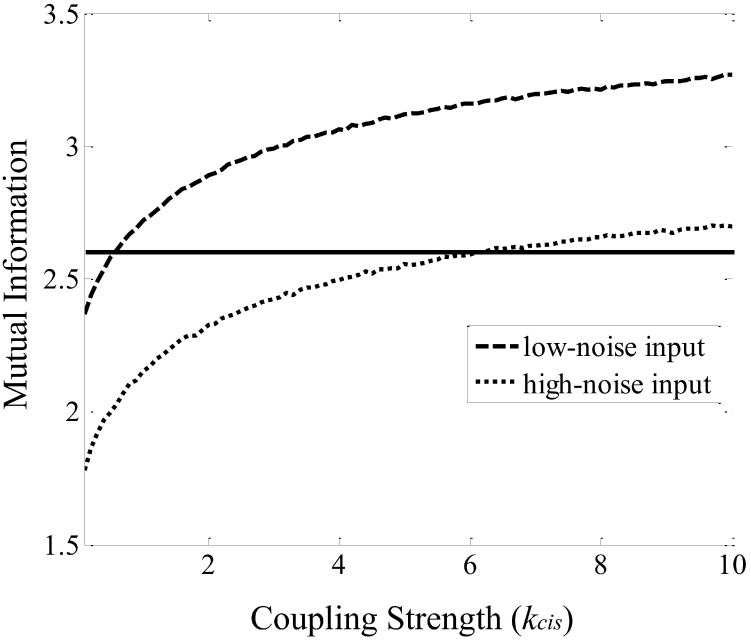
Mutual information of the Notch-Delta channel with respect to the coupling strength for low noise, *σ*^2^ = 0.01 (dashed line) and high noise, *σ*^2^ = 0.03 (dotted line). The horizontal solid line depicts the same level of the mutual information between input-output which can be achieved by changing the mutual coupling of Notch-Delta. The input *D^ext^* is normally distributed around *μ* = 1 and the parameter values are: *β*_N_ = 1, *β_D_* = 100, *γN =γD* = 1, *k_+_* = 2, *k_*= 1, Γ = 100.

**FIG. 3.**
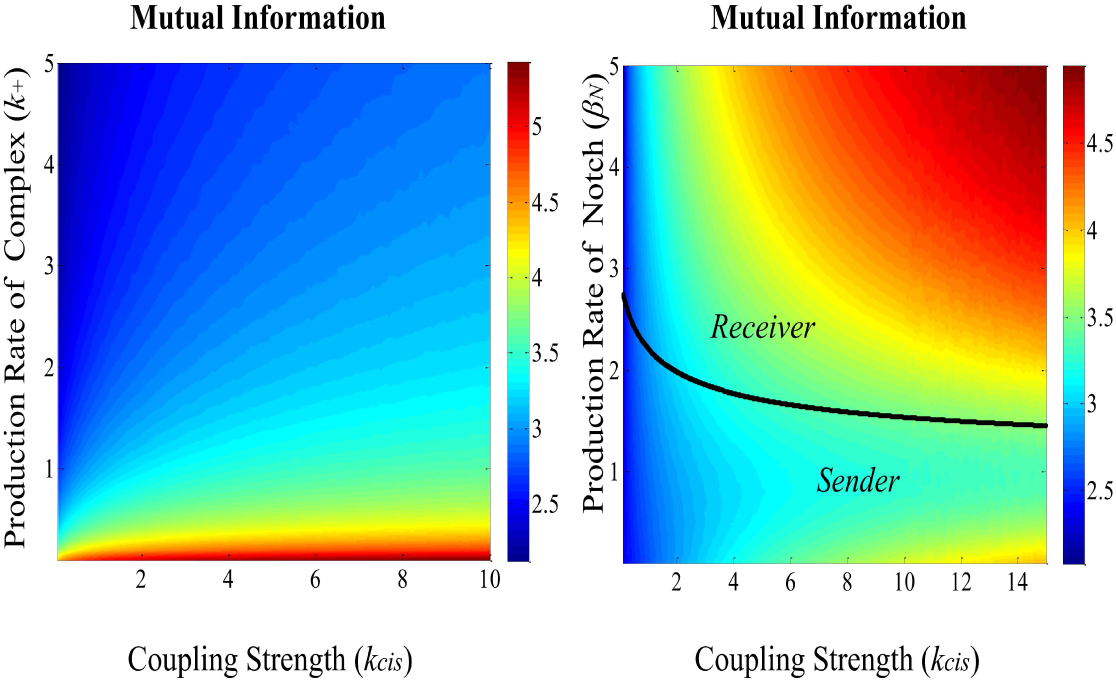
Notch-Delta channel performance for different parameters. (a) Mutual Information dependency on the coupling strength (k_cis_) and the production rate of complex (*k_+_*). The input *D^ext^* is normally distributed around *μ* = 1 with *σ*^2^ = 0.01 and the parameter values are: *β_N_* = 1, *β_D_* = 100, *γN* = *γD* = 1, *k*_ − = 1, Γ = 100. (b) Mutual Information dependency of the coupling strength (*k_cis_*) and the production rate Notch (*β_N_*). The solid line depicts the critical point that the cell switches state between sender and receiver. The input *D^ext^* is normally distributed around *μ* = 1 with *σ*^2^ = 0.01 and the parameter values are: *β_D_* = 100, *γN* = *γD* = 1, *k_+_* = 2, *k_* = 1, Γ = 100.

In the following, we investigate how the system’s performance depends on the production rates of Notch *βN* and Complex *k*_+_ and the coupling strength of Notch-Delta *k_cis_*. By increasing the cell’s encoding strength *k*_+_ the input-output mutual information decreases. On the other hand, as shown in our simple communication model, increasing the coupling strength *k_cis_* always enhances the fidelity of the Notch-Delta channel (Figure 3(a)). This result indicates that the rise in intracellular coupling strength can compensate for the loss of the cell’s communication performance induced by increasing the encoding rate.

**FIG. 4.**
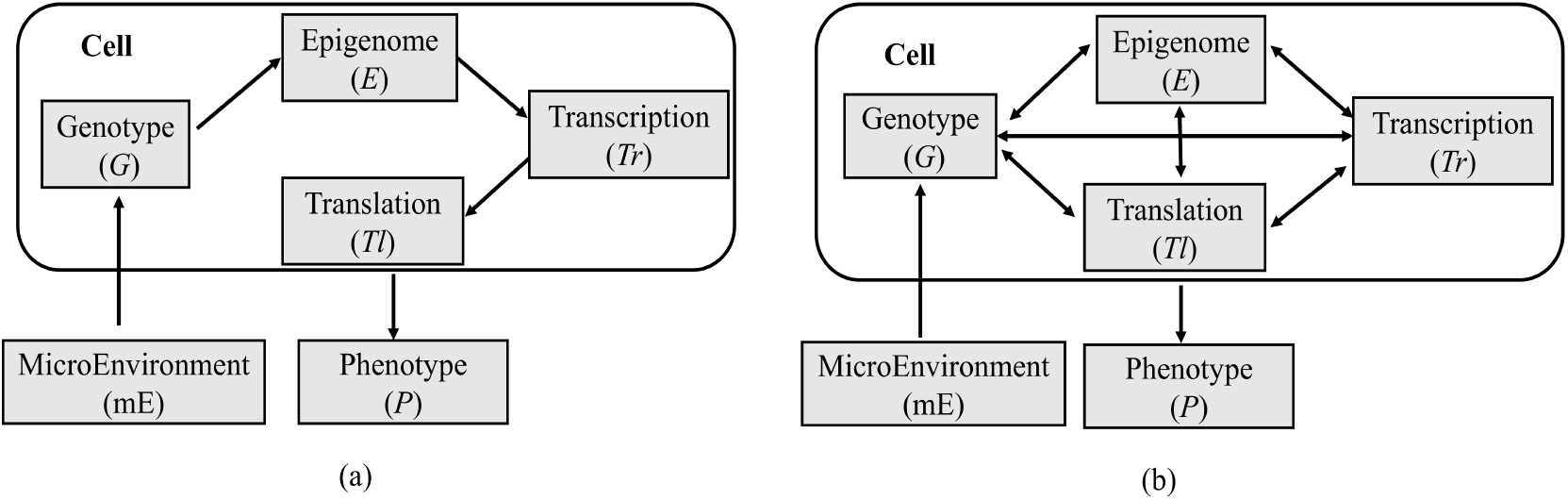
Cellular encoding. Hypothesis A: serial processing of intracellular information. (b) Hypothesis B: parallel processing of intracellular information.

By variation of both the production rate of Notch *β_N_* and the coupling strength *k_cis_* we can identify the two cell states with respect to the Notch-Delta pathway: the receiver state, where the equilibrium of Notch is greater than the corresponding Delta (*N_eq_ > D_eq_*) and the sender state where (*N*_*eq*_ < *D*_*eq*_) (Figure 3(b)). Please note that the mixed state (where *N*_*eq*_ ≈ *D*_*eq*_) is identified with a precursor state where the cells are not differentiated yet [10]. While increasing the coupling strength *k*_*cis*_ always improves the performance of the Notch-Delta channel, the same is not true with respect to the production rate of Notch. Close to the critical point, which marks the transition from sender to receiver, the Notch-Delta channel has a poor performance. The latter information-theoretic result is in agreement with previous results indicating the dynamical behaviour of differentiating cells when they undergo a critical state transition before fate selection [15].

The cell in a receiver state increases its performance by increasing Notch production rate *β_N_*. The cell in a sender state increases its performance by reducing *β_N_*. This implies that the cell should stop investing energy in producing Notch receptors when it finds itself in a sender state. We can conjecture that improving performance leads to an emergent energy management mechanism. This could experimentally be tested by probing cells “locked” in a sender state for different nutrient/energy conditions and measuring the corresponding Notch production rate.

## Coupling in cell encoding

Our postulated principle that intracellular coupling improves the cell decision-making reliability has more fundamental implications. Typically, the cell encodes environmental cues (*mE*) into the genetic (*G*), epigenetic (*E*), transcriptional (*Tr*) and translational (*Tl*) levels. The decoded output is being represented by the resulting phenotype (*P*). Genetic textbook knowledge dictates that phenotypic responses result from the serial/hierarchical processing of the above mechanisms, as *mE → G → E → Tr → Tl → P* (Hypothesis A) [16]. Recently, an alternative hypothesis has been postulated that assumes all intracellular processing levels are interconnected implying a parallel cellular encoding (Hypothesis B) [16]. Assuming that each processing level is a channel, we can draw two different communication theoretic scenarios as in Fig. 4.

If we assume that Hypothesis A is true, we can state that, using the data processing theorem, the serial processing implies a loss of information, since *I*(*mE, P*) ≤ *I*(*mE, X*_*i*_) where *X_i_* = *G, E, Tr, Tl*. If Hypothesis B is true, it implies that a cell processing not only preserves input information, but also produces even more (in analogy to Eq. (5) and [7]). The excess information produced could be transmitted back to the cellular microenvironment [4]. Therefore, by the virtue of improved phenotypic response fidelity, Hypothesis B of coupled encoding levels is more likely to be biologically relevant. Finally, using the concept of directed information one can identify the information flow among intracellular encoding levels [17].

## Conclusions

We firmly believe that intracellular coupling is a salient feature of cell decision-making. As shown above, it confers an advantage to cells since it increases their ability to process noisy microenvironmental information. We postulate that intracellular network coupling is a structural design principle that allows cells to generate reliable responses in noisy environments. Additionally, as shown for the Notch-Delta pathway, optimising cell responses can be associated with an energy management mechanism. In the future, we need to experimentally test our postulated principle and its implications for further signal transduction pathways and confirm its generality.

AIR and HH gratefully thank Prof. M. Meyer-Hermann, Prof. C. Siettos and Prof. T. Krueger for their valuable comments and fruitful discussions. HH and AIR would like to acknowledge the SYSMIFTA ER-ACoSysMed grant (031L0085B) for the financial support of this work.

